# Spontaneous enteric nervous system activity precedes maturation of gastrointestinal motility

**DOI:** 10.1101/2023.08.03.551847

**Authors:** Lori B. Dershowitz, Hassler Bueno Garcia, Andrew S. Perley, Todd P. Coleman, Julia A. Kaltschmidt

## Abstract

Spontaneous neuronal network activity is essential in development of central and peripheral circuits, yet whether this is a feature of enteric nervous system development has yet to be established. Using *ex vivo* gastrointestinal (GI) motility assays with unbiased computational analyses, we identify a previously unknown pattern of spontaneous neurogenic GI motility. We further show that this motility is driven by cholinergic signaling, which may inform GI pharmacology for preterm patients.

## Introduction

In many regions of the embryonic nervous system, neural circuit activity develops before the experience of sensory cues.^1^ This spontaneous neuronal network activity facilitates functional circuit maturation.^2^ A quintessential example of spontaneous activity is retinal waves, the intermittent bursting and silent periods of retinal ganglion cells that arise prior to visual stimuli.^1,3^ In the developing spinal cord, spontaneous bursting activity is required for proper motor axon finding.^2,4^ Whether similar spontaneous activity occurs in the developing enteric nervous system (ENS), the largest branch of the peripheral nervous system, remains to be explored.

The ENS resides in the walls of the intestine and autonomously controls gastrointestinal (GI) functions including GI motility. Development of the GI motility circuit within the ENS is necessary for the propulsion of intestinal matter. Failure in ENS development underlies pediatric disorders such as Hirschsprung’s disease, in which a segment of intestine develops without ENS innervation and cannot pass fecal matter. Despite the clinical need to understand maturation of the ENS circuit that controls GI motility, few studies have investigated the earliest events in the development of this circuitry.

The first form of GI motility is non-neurogenic. Myogenic ripples, bidirectional GI motility that arises from the intestinal smooth muscle, appear prior to neurogenic GI motility in *ex vivo* analyses of both embryonic mouse^5^ and human fetal intestines.^6^ Mouse myogenic ripples were first observed at embryonic day (E) 13.5 in small intestine (SI) and at E14.5 in colon.^5^ Human myogenic ripples have been observed as early as 14 postconceptional weeks.^6^ In both embryonic mouse and human fetal intestines, this motility pattern is unperturbed by tetrodotoxin (TTX), a sodium channel blocker used to silence neuronal activity.^5,6^ Another non-neurogenic form of GI motility is slow waves, rhythmic contractions of smooth muscle controlled by interstitial cells of Cajal.^7^ Although the exact age of slow wave development is not known, electrical recordings of intestinal smooth muscle in mouse have identified slow waves at postnatal day (P) 0 in SI and P10 in colon.^8^

Neurogenic GI motility arises in the late embryonic and early postnatal period in mice. While activity of individual enteric neurons has been observed as early as E11.5, neurogenic GI motility has not been observed until E18.5.^5,9^ Roberts et al. identified the first distally propagating contractions by monitoring *ex vivo* motility of the mouse embryonic SI and colon.^5,10^ They detected contraction complexes in the E18.5 duodenum^5^ and found colonic migrating complexes in the P10 colon.^10^ However, the earliest forms of activity in the embryonic retina and spinal cord do not resemble mature circuit function,^2^ posing the question whether earlier neurogenic motility signatures can be detected in the developing gut.

Here, we utilized an *ex vivo* embryonic whole-intestine motility monitor and developed an unbiased analytical approach to identify a pattern of neurogenic GI motility termed “clustered ripples” that is distinct from adult GI motility and precedes all known forms of neurogenic GI motility. Clustered ripples consists of dynamic clustering of myogenic ripples that travel the length of the SI and exhibits intermittent on/off periods, reminiscent of early spontaneous activity in the retina and spinal cord.^2^ We use pharmacologic and genetic approaches to demonstrate that this pattern requires ENS synaptic activity and that cholinergic and nitrergic signaling play opposite roles in clustering. Collectively, our work unveils patterned neurogenic GI motility that occurs in the absence of external stimuli and precedes the earliest forms of neurogenic GI motility in mice.

## Results

To visually assess development of GI motility, we dissected whole mouse intestines from E14.5-E18.5 embryos and placed the intestines in an organ bath with warmed circulating Krebs solution and an attached camera, a setup called an *ex vivo* GI motility monitor (GIMM).^11,12^ We surveyed video recordings for evidence of GI motility that could be visually detected (Fig. 1A-C and supplementary videos). Given that both myogenic and neurogenic GI motility arise in the SI prior to the colon,^5,10^ we focused on the SI for our analyses. At E14.5, little GI motility was noted (Fig. 1A and supplementary videos). At E16.5, we observed isolated “jerking” movements (Fig. 1B, asterisk, and supplementary videos), and at E18.5 GI motility appeared more frequent and complex (Fig. 1C).

**Figure 1:**
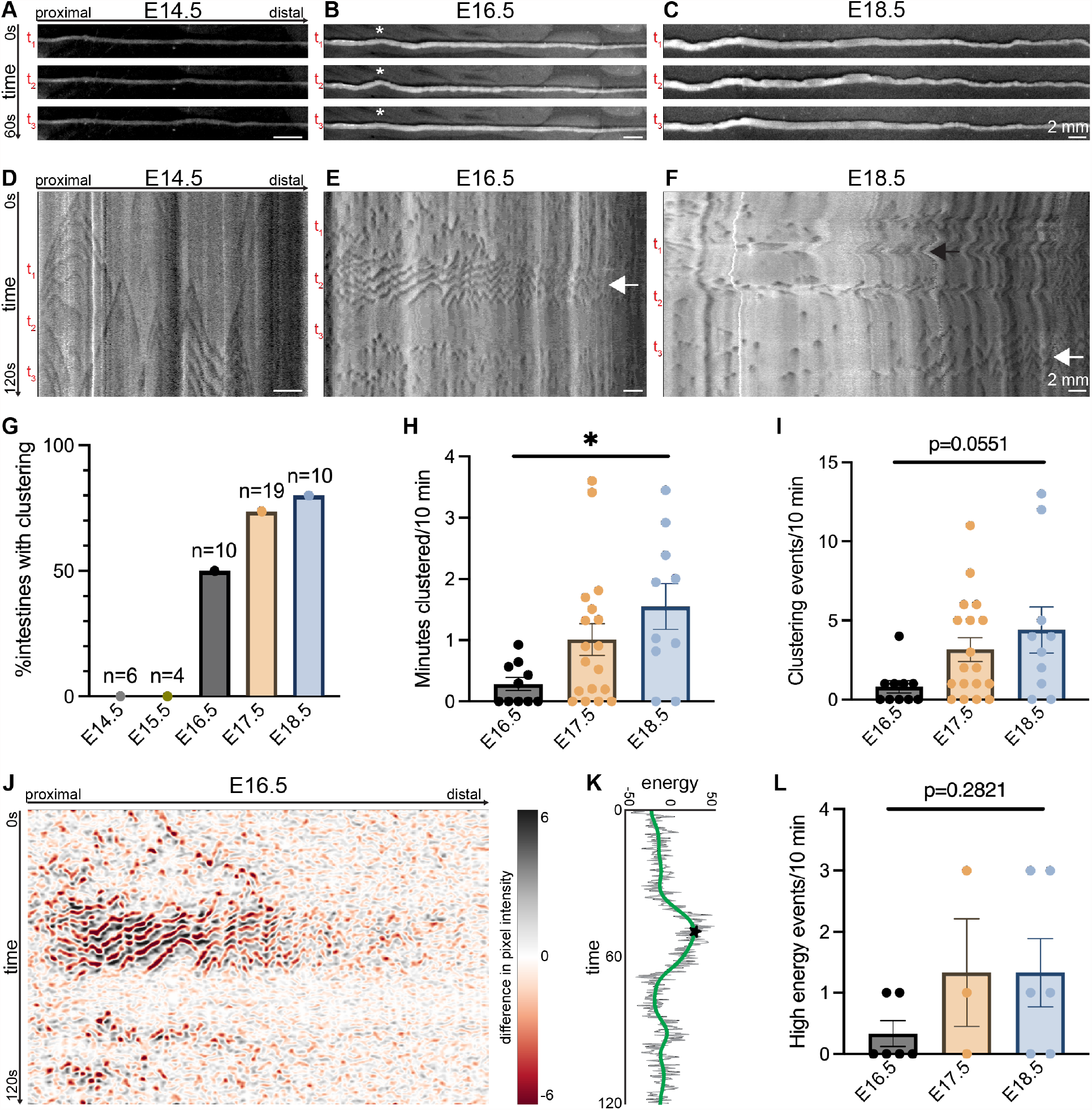
Clustered ripples arise at E16.5 in the mouse small intestine. (**A-C**) Representative still images spanning 60 seconds of video captured in the *ex vivo* gastrointestinal motility monitor (GIMM) from embryonic day (E)14.5 (A), E16.5 (B), and E18.5 (C) mouse small intestine (SI). Asterisk indicates jerking motility in the E16.5 intestine. Scale bars as indicated. (**D-F**) Representative spatiotemporal maps (STMs) of *ex vivo* GI motility in the E14.5 (D), E16.5 (E), and E18.5 (F) mouse SI. Dark gray: decreased intestinal diameter; light gray: increased. t_1_-t_3_denote timepoints of still images in A-C. White arrows indicate clustered ripples; black arrow indicates a contraction complex. Scale bars as indicated. (**G**) Percentage of intestines from E14.5-E18.5 exhibiting clustered ripples during a 10-minute window in the *ex vivo* GIMM. n per given age as indicated in graph. (**H**) Total minutes exhibiting clustering during a 10-minute window from E16.5, E17.5, and E18.5 SI. n=10-19 intestines per age. \**p*<0.05 from one-way ANOVA. (**I**) Number of individual clustering events observed in a 10-minute window from E16.5, E17.5, and E18.5 SI. n=10-19 intestines per age. *p*-value from one-way ANOVA as indicated on graph. (**J**) Representative postprocessed plots of E16.5 STM in E. Black and red: large changes in pixel values over time; white: no change in pixel value. (**K**) Representative energy plot derived from the postprocessed STM in (J). Gray line denotes the energy time series. Green line denotes the smoothed output. Star indicates high energy event. (**L**) Number of high energy events identified in a 10-minute window from E16.5, E17.5, and E18.5 SI. n=3-6 intestines per age. *p*-value from one-way ANOVA as indicated on graph. Pairwise comparisons in Supplementary Table 1.

For a more granular analysis of the observed GI motility patterns, we generated spatiotemporal maps (STMs) of *ex vivo* GI motility in the developing mouse SI. STMs graphically describe changes in intestinal diameter over time with darker shades denoting contractions and lighter shades denoting relaxation.^13^ At E14.5, STMs reveal bidirectional GI motility events that correspond to previously described myogenic ripples (Fig. 1D). At E16.5, myogenic ripples persist, yet they are periodically clustered together along the length of the SI (Fig. 1E, white arrow). This GI motility pattern, which we termed “clustered ripples,” occurs at timepoints that correspond to the observed jerking motility (Fig. 1B, E). Clustered ripples do not have a directional bias and are followed by a period of quiescence in which the contour of the intestine is smooth and STMs do not indicate any motility (Fig. 1B, E).

We next assessed clustered ripples in STMs over developmental time. Clustered ripples are not observed at E14.5 or at E15.5. The percentage of embryonic intestines exhibiting clustering increases from 50% at E16.5 to 80% at E18.5 (Fig. 1G). To quantify characteristics of clustered ripples, we manually evaluated the total time each sample exhibited clustering as well as the number of clustering events during a 10-minute acquisition. The overall time spent in clustered ripples (∼0.3 min at E16.5, ∼1.0 min at E17.5, and ∼1.6 min at E18.5) and number of clustering events (∼0.8 at E16.5, ∼3.2 at E17.5, and ∼4.4 at E18.5) increased over developmental time (Fig. 1H, I). At E18.5, we still observe clustered ripples though they tend to be isolated to the distal portion of the SI (Fig. 1F, white arrow). At this age we also begin to observe previously described contraction complexes (Fig. 1F, black arrow).

To confirm that clustered ripples represent a distinct motility phenomenon, we generated an unbiased computational method to analyze STMs. By assessing how pixel intensity changes over time, this approach identifies “high energy” points in time, or moments with increased probability of observing a motility event compared to baseline (Fig. 1J, K). These high energy events correspond to clustered ripples. When applied over multiple ages, this approach identified a similar trend in clustered ripple development as we found with manual quantification (Fig. 1L).

To determine whether clustered ripples are a neurogenic form of GI motility, we next assessed *ex vivo* GI motility after silencing neurons. We silenced activity of enteric neurons pharmacologically by adding nonspecific sodium channel blocker tetrodotoxin (TTX) to the *ex vivo* GIMM bath (Fig. 2A-D). TTX broadly inhibits action potentials and has been previously used to silence enteric neurons in embryonic and adult mice.^5,14^ In parallel, we used *Wnt1-Cre*;*R26*^floxstop-TeNT^ mice to selectively express tetanus toxin light chain subunit (TeNT) in neural crest derivatives, which include only the ENS in intestinal tissue. Expression of TeNT inhibits synaptic vesicle fusion and silences synaptic transmission. As anticipated, both pharmacologically and genetically silencing neurons does not alter myogenic ripples at E14.5 as TTX and TeNT do not inhibit other mediators of GI motility including smooth muscle and interstitial cells of Cajal (Fig. A, E). However, both of these approaches completely ablate clustering activity in all intestines assayed from ages E16.5-E18.5 (Fig. 2C, D, J, K). We used the computational approach to confirm that high energy events are absent when enteric neurons are silenced (Fig. 2E-G). Taken together, these results demonstrate that enteric neuron activity is required to generate clustered ripples. Other mediators of GI motility, including interstitial cells of Cajal and intestinal smooth muscle, are not inhibited by TTX and are therefore not responsible for clustering.

**Figure 2:**
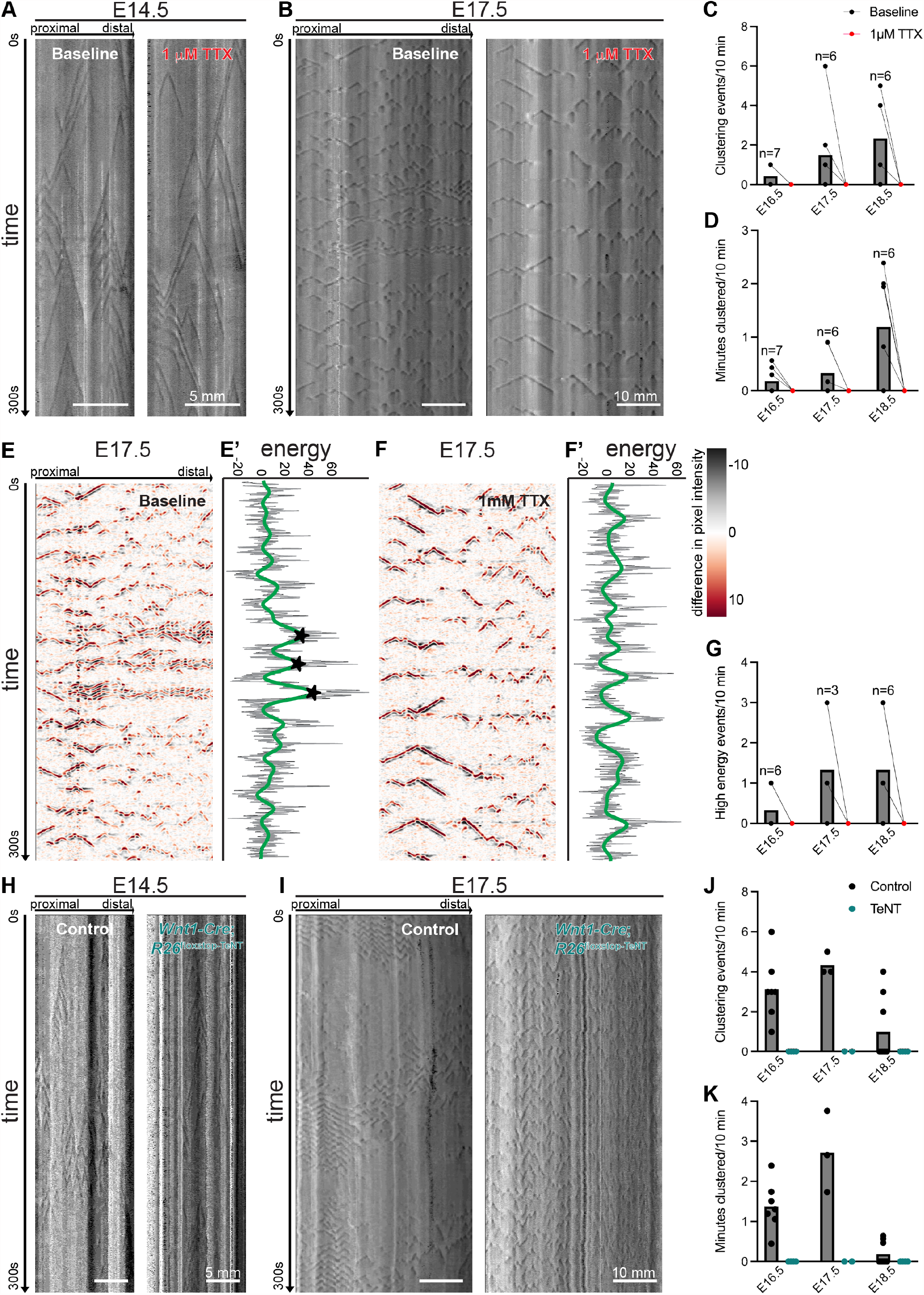
Clustered ripples are a neurogenic form of GI motility. (**A**) Representative STMs of *ex vivo* GI motility in the E14.5 SI at baseline (left) and after the addition of 1 mM tetrodotoxin (TTX, right). Dark gray: decreased diameter; light gray: increased. Scale bars as indicated. (**B**) Representative STMs from the E17.5 SI at baseline (left) and with 1 mM TTX (right). Dark gray: decreased diameter; light gray: increased. Scale bars as indicated. postprocessed plots of STMs. Black and red: large changes in pixel values over time; white indicates: no change in pixel value. (**C**,**D**) Number of clustering events (C) and minutes clustered (D) during a 10-minute window at baseline (black) and after addition of 1 mM TTX (red) in the E16.5, E17.5, and E18.5 SI. n as indicated on graph per age. (**E-F’**) Postprocessed plots of STMs in (B) with accompanying energy plots. Black and red: large changes in pixel values over time; white: no change in pixel value. Gray line denotes the energy time series. Green line denotes the smoothed output. Stars indicate high energy events identified through the unbiased approach. (**G**) Quantification of number of high energy events identified in a 10-minute window at baseline (black) and after addition of 1 mM TTX (red) from E16.5, E17.5, and E18.5 SI. n as indicated on graph per age. (**H**) Representative STMs of *ex vivo* GI motility in *Cre* negative control (left) and *Wnt1-Cre*;*R26*^floxstop-TeNT^ (right) at E14.5. Dark gray: decreased diameter; light gray: increased. Scale bars as indicated. (**I**) Representative STMs from a *Cre* negative control (left) and *Wnt1-Cre*;*R26*^floxstop-TeNT^ (right) SI at E17.5. Dark gray: decreased diameter; light gray: increased. Scale bars as indicated. (**J**,**K**) Number of clustering events (I) and minutes clustered (J) during a 10-minute window in *Cre* negative controls (black) and *Wnt1-Cre*;*R26*^floxstop-TeNT^ mice (teal) at E16.5, E17.5, and E18.5. n=2-9 per genotype per age as indicated. Pairwise comparisons in Supplementary Table 1.

Cholinergic excitatory motor neurons and nitrergic inhibitory motor neurons are the main drivers of GI motility within the ENS, and these two neuronal populations play distinct roles in adult mouse GI motility.^15^ To determine whether cholinergic and nitrergic enteric neurons play a role in clustered ripples, we pharmacologically inhibited either subclass in the *ex vivo* GIMM preparation. For this analysis we focused on E17.5, an age at which 73% of embryonic intestines exhibit clustered ripples but contraction complexes are not yet detected (Fig. 1G).^5^ To inhibit cholinergic signaling, we added the nicotinic antagonist mecamylamine, which has been used to inhibit cholinergic signaling in the adult guinea pig colon and to silence spontaneous activity in the embryonic mouse spinal cord.^4,16^ At E17.5, addition of mecamylamine to the GIMM bath completely quenched clustered ripples in all intestines examined (Fig. 3A-C). To inhibit nitrergic signaling, we used N_ω_-nitro-L-arginine (NOLA). NOLA has previously been used to silence nitrergic signaling in the neonatal mouse colon.^10^ Addition of NOLA to the GIMM bath increased both the number of clustering events and the minutes clustered (Fig. 3D-F). The effect was often dramatic with large periods of clustering and quiescence (Fig. 3D). Thus, cholinergic signaling is required for generating clustered ripples in the embryonic mouse SI, and nitrergic signaling appears to have an opposing effect and reduce clustering in the embryonic mouse SI. Collectively, these results suggest that cholinergic and nitrergic signaling play opposite roles in the generation of clustered ripples.

**Figure 3:**
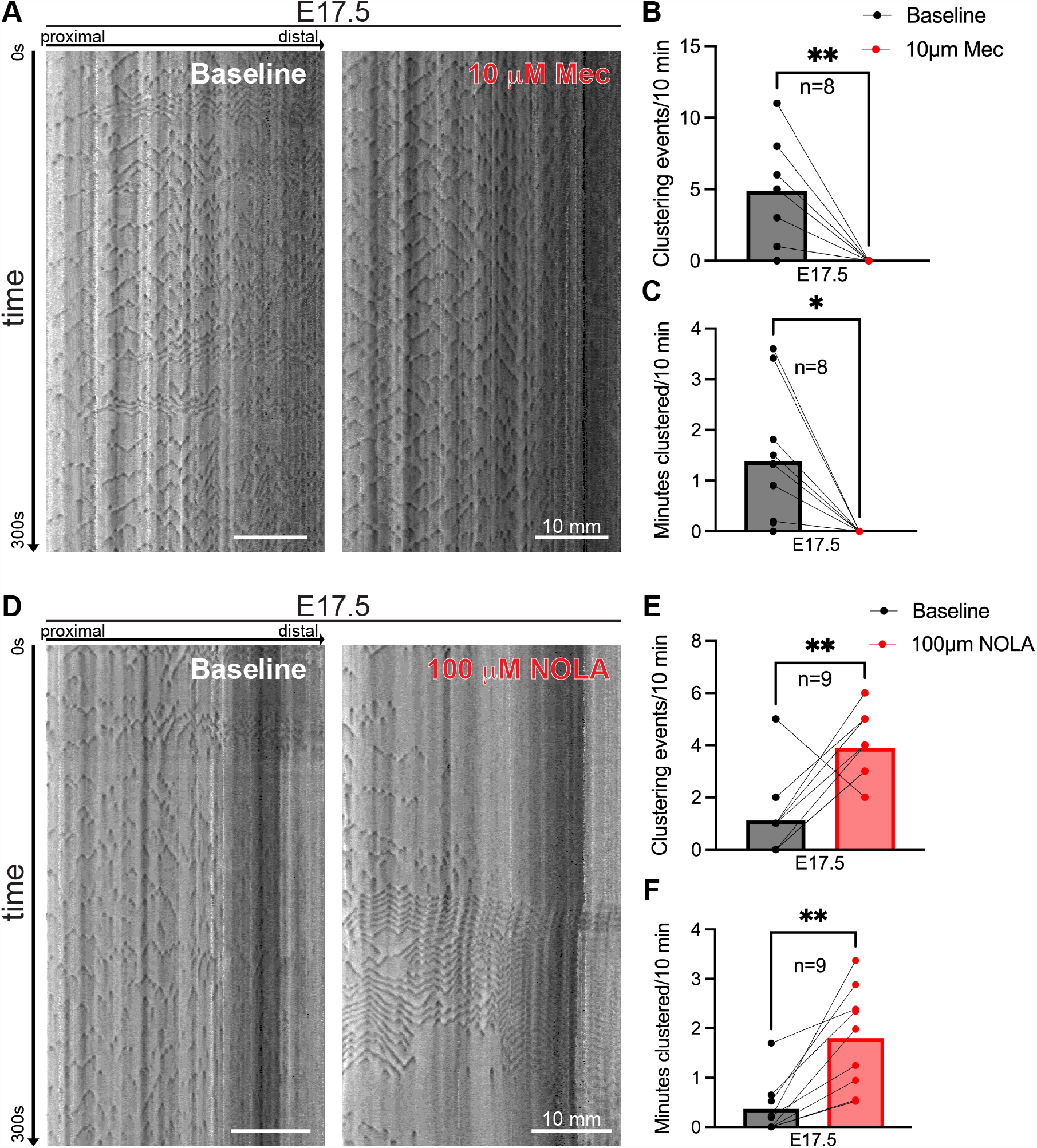
Cholinergic and nitrergic signaling have opposing roles in clustered ripples. (**A**) Representative STMs of *ex vivo* GI motility in the E17.5 SI at baseline (left) and after the addition of 10 mM mecamylamine (Mec, right). Dark gray: decreased intestinal diameter; light gray: increased. Scale bars as indicated. (**B**,**C**) Number of clustering events (B) and minutes clustered (C) during a 10-minute window at baseline (black) and after addition of 10 mM Mec (red) at E17.5. n=8. (**D**) Representative STMs from the E17.5 SI at baseline (left) and after the addition of 100 mM N_ω_-nitro-L-arginine (NOLA, right). Dark gray: decreased diameter; light gray: increased. Scale bars as indicated. (**E**,**F**) Number of clustering events (E) and minutes clustered (F) during a 10-minute window at baseline (black) and after addition of 100 mM NOLA (red) at E17.5. n=9. All tests paired student t-tests. \**p*<0.05, \*\**p*<0.01.

## Discussion

Here we identified and characterized clustered ripples, a form of spontaneous ENS activity that arises prior to any known form of neurogenic GI motility. We first observed clustered ripples in the E16.5 SI, and the instances and length of clustering increase over developmental time from E16.5-E18.5. We found that activity of enteric neurons is necessary to generate clustering, yet different enteric neuron subclasses play distinct roles with cholinergic neurons enhancing clustering and nitrergic neurons opposing clustering. We further present an unbiased approach to identify motility events in STMs, which allowed us to computationally and quantitatively describe clustered ripples and confirm their absence when enteric neuron activity is inhibited.

In both the developing retina and cerebellum, spontaneous activity propagates over long distances similar to how clustered ripples can span the length of the SI.^2^ GI motility patterns that span long distances have been previously described. In adult mice, a form of neurogenic GI motility called motor complexes can travel long distances and cross the ileocecal junction.^17^ In contrast, embryonic GI motility such as contraction complexes are discrete. Previous *ex vivo* GI motility studies of the embryonic SI cut the tissue at the ileocecal junction and did not observe clustered ripples.^5^ Therefore, we suggest that continuity of the SI, cecum, and colon may be necessary for observing clustered ripples in *ex vivo* preparations.

Another shared feature of spontaneous neurogenic activity in the developing retina, spinal cord, hippocampus, cochlea, and cerebellum is bursting behavior.^2^ This is akin to the on/off behavior we observe with clustered ripples, which arise across the length of the SI followed by a period of global motility quiescence. Given that clustered ripples exhibit on/off behavior across a large expanse of the GI motility circuit, clustered ripples may represent a parallel developmental phenomenon to spontaneous neural network activity in other regions of the developing nervous system.

In this study, we utilize the autonomous nature of the ENS and *ex vivo* GI motility as a readout of neuronal activity. One limitation of the *ex vivo* GI motility readout is that it requires ENS innervation of the smooth muscle. In the developing chick spinal cord, spontaneous activity of motor neurons precedes their innervation of target muscles.^18^ The exact age at which enteric neurons innervate the intestinal smooth muscle is not known, yet histological analyses have visualized nerve terminals in the intestinal smooth muscle as early as E17.5.^19^ To assess for spontaneous ENS activity that precedes innervation, future studies should utilize approaches such as calcium imaging and electrophysiology to assess enteric neuron network activity at E15.5, an age just prior to the emergence of clustered ripples.

What is the role of clustered ripples in development of the ENS and GI motility? As mentioned above, clustered ripples do not have a directional bias and are thus unlikely to play a role in propulsion of intestinal contents. Instead, clustered ripples may play a transitory role in establishment and refinement of the circuitry underlying GI motility. In the adult ENS, dynamic interplay between cholinergic and nitrergic signaling underlies motor complexes. Cholinergic neurons exhibit increased activity during motor complexes and decreased activity between GI motility events; nitrergic neurons exhibit the opposite firing tendencies.^15^ Our data suggest that these two neuronal populations also act in opposition developmentally (Fig. 3.3A-F). Prior studies have shown that enteric neurons destined to become adult cholinergic or nitrergic neurons are born in the mid-to-late embryonic period in mice, around the time when clustered ripples arise.^19^ Therefore, clustered ripples could potentially represent a developmental tug-of-war establishing the balance of excitation and inhibition required for GI motility.

Clustered ripples are the earliest identified form of spontaneous enteric nervous system activity and may serve vital roles entraining the circuitry for GI motility. Spontaneous network activity is required for circuit development in tissues such as in the visual system and spinal cord.^2^ Whether clustered ripples are necessary for GI motility circuit development remains to be explored. The relevance of this question extends to human gut development, as we have observed clustered ripples in the human fetal SI.^6^ Currently, there are no drugs to enhance GI motility in extremely preterm infants, which are developmentally similar to 2^nd^ and 3^rd^ trimester fetuses. The human fetal ENS has increased representation of nitrergic neurons relative to cholinergic neurons.^6^ Here, we show that inhibiting nitrergic signaling increases clustered ripples, an observation which may suggest potential pharmacologic targets to enhance GI motility and increase survival in the preterm population. Therefore, understanding the physiology of clustered ripples may provide fundamental insights into the development of GI motility, a function critical to survival.

## Methods

### Mice

All procedures followed the National Institutes of Health Guidelines for the Care and Use of Laboratory Animals and were approved by the Stanford University Administrative Panel on Laboratory Animal Care. Adult mice were housed up to five per cage. Mice were given food and water *ad libitum* and were maintained on a 12 hour/12 hour light/dark cycle. In order to date embryonic age, female mice were visually inspected for a vaginal plug each morning, and plugged females were separated from the breeding cage. Embryos were collected from ages spanning embryonic day (E) 14.5-E18.5.

Unless otherwise specified, experiments were performed on C57BL/6J mice (The Jackson Laboratory). Other mouse lines in this study include *Wnt1-Cre* (Gift of Becker lab, Stanford University School of Medicine)^20^ and *R26*^floxstop-TeNT^ (Gift of Goulding Lab, Salk Institute).^21^ Both *Wnt1-Cre* and *R26*^floxstop-TeNT^ mice were maintained on a mixed background. Controls for experiments with *Wnt1-Cre*;*R26*^floxstop-TeNT^ mice were *Cre* negative littermate embryos.

### Ex vivo *motility analysis*

The *ex vivo* GIMM setup and analysis was adapted from previous studies.^11–13^ Pregnant adult females were culled with CO_2_and cervical dislocation. Embryos were immediately removed and placed in 37 °C Krebs solution (pH 7.4 containing (in mmol/l): 117 NaCl, 4.7 KCl, 3.2 CaCl_2_ (2H_2_O), 1.5 MgCl_2_(6H_2_O), 25 NaHCO_3_, 1.2 NaH_2_PO_4_and 11 Glucose). For experiments with *Wnt1-Cre*;*R26*^floxstop-TeNT^ embryos, tails were collected at this moment for subsequent genotyping. While submerged in warmed Krebs, embryonic intestines were dissected from stomach to anus under a dissecting microscope, and the mesentery was gently cut in order to straighten the intestine. Intestines were immediately moved to an organ bath with circulating 37 °C Krebs solution and a heated water bath below to maintain chamber temperature. The intestines were tautly pinned at the stomach and anus on Sylgard 170 in the chamber base. Intestines were allowed to acclimate in the organ bath undisturbed for 10 minutes. A high-resolution monochromatic firewire industrial camera (Imaging Source®, DMK41AF02) connected to a 2/3” 16 mm f/1.4 C-Mount Fixed Focal Lens (Fujinon HF16SA1) was mounted above the organ bath, and 10-minute videos were captured at 3.75 frames/second using IC capture software (Imaging Source). 30 minutes of videos were captured at baseline. For experiments involving drugs, an additional 30 minutes were recorded with the drug followed by 30 minutes of washout.

STMs were generated from video recordings with VolumetryG9a software.^13^ For each experiment, the final 10-minute video per condition was analyzed. STMs were visually inspected for GI motility patterns including myogenic ripples, clustered ripples, and contraction complexes. The number of contraction events per 10-minute STM was manually counted, and the minutes clustered were counted using the line segment and measure functions in ImageJ/FIJI.

### Quantitative analysis of spatiotemporal dynamics

Image processing was done through algorithms available in SciPy^22^ and consisted of an outlier detection to remove noisy measurements, filters to improve the signal to noise ratio, and a peak detection algorithm over time to identify epochs of clustered ripples. The spatiotemporal image experienced artifacts that were identified via outlier detection. Since the ripples (contractions) presented as sharp changes in pixel intensity, an edge detection filter, commonly used in image processing, was applied to identify the ripples. Specifically, a Sobel filter was applied across the time axis to highlight these changes in pixel intensity over time.^23^ The spatiotemporal output was then collapsed in space using a weighted sum, where the weights correspond to the inverse of the variance observed across each spatial measurement. The temporal output was then processed using a peak finding algorithm to detect and filter peaks associated with clustered ripples.

The specifics of the methods are as follows: the spatiotemporal images were originally stored as a Tag Image File Format (TIFF) and imported to Python using the integrated image processing library Pillow. With this library, the images were converted to greyscale and copied into a NumPy array, for signal processing with SciPy. The data cleaning procedure consisted of identifying artifacts in the image that presented themselves as (i) salt and pepper noise due to bubbles in the recording, and (ii) artifact lines in time due to spatial locations with too narrow a diameter to be recognized by VolumetryG9a software.

To detect salt and pepper noise, each pixel was classified as artifact based on a thresholding range computed on the statistics of the measured signal, specifically the median plus or minus 4 standard deviations. Once these pixels were identified, neighboring samples were also identified as outliers keeping in mind that these outlier pixels would affect the output of future processing steps. Additionally, artifacts in time were detected by going through each spatial vector in time and identifying completely saturated measurements (e.g. the standard deviation of one time location across space was 0). The identification of outliers was then used to minimize their effect on the processed output.

To identify clustered ripples, it was noted that they presented themselves as sharp changes in pixel intensity and a Sobel filter (scipy.ndimage.sobel) was applied across the time domain to highlight pixel intensity over time. Additionally, pixels that were previously identified to be salt and pepper outliers were replaced by zero values at this stage. Furthermore, the spatial axis was collapsed by computing the weighted sum of all spatial values at one time point (scipy.spatial.distance.euclidean), where each weight was the inverse of the variance across one spatial location. For time epochs that were identified as outliers, their value was replaced by the median of the time series. A Gaussian filter (package: scipy.ndimage.gaussian_filter1d) was subsequently applied to smooth the temporal output, where the parameter sigma was chosen to be 15 sample points. Lastly, to identify ripples, we identified local maxima in the output of the Gaussian filter by implementing a peak-finding algorithm (scipy.signal.find_peaks with height parameter determined by 1 standard deviation above the mean of the time series computed before Gaussian smoothing and width parameter given by 20 sample points).

### Drugs

Drugs were stored as aliquots diluted in distilled H_2_O with the exception of N_ω_-nitro-L-arginine, which was diluted in a solution hydrochloric acid titrated with H_2_O to pH 1.8. To analyze the effects of inhibiting all enteric neurons or subpopulations thereof, drug aliquots were diluted to desired concentration in 37 °C Krebs solution, and the drug-containing Krebs solution was circulated through the organ bath. GI motility was analyzed after 20 minutes of a given drug circulating through the organ bath. The following drugs and concentrations were used: 1 mM tetrodotoxin (Alomone Labs), 10 mM mecamylamine (Sigma-Aldrich), and 100 mM N_ω_-nitro-L-arginine. Concentrations were selected based on prior studies using these drugs in the ENS and in the embryonic mouse nervous system.^5,10,16,24^

### Graphing and statistical analyses

All graphing and statistical analyses were performed with Prism 9 software (GraphPad). Statistical analysis with one-way ANOVA was performed to determine if age was a significant factor in number of events and length of clustering. Paired student t-tests were used to compare means of events and length of clustering between baseline and addition of a drug. Unpaired student t-tests were used to compare means of events and length of clustering between *Wnt1-Cre*;*R26*^floxstop-TeNT^ mice and *Cre* negative littermate embryos.

## Author Contributions

L.B.D designed, performed, and analyzed all experiments and co-wrote the manuscript. H.B.G., A.S.P., and T.P.C. designed and implemented computational analyses. J.A.K. designed and supervised experiments and co-wrote the manuscript.

## Acknowledgements

We would like to thank the entire Kaltschmidt lab, especially Jennifer Lee Shadrach, for feedback on experimental design and results. We would also like to thank Dr. Laren Becker for supplying the *Wnt1-Cre* mice and Dr. Martyn Goulding for supplying the *R26*^floxstop-TeNT^ mice. This work was supported by Stanford Medical Scientist Training Program grants T32 GM007365-44 and T32-GM145402 (L.B.D.), the Stanford Data Science Scholars Program (A.S.P), the Wu Tsai Neurosciences Institute (T.P.C. and J.A.K.), the Stanford University Department of Neurosurgery, and research grants from The Shurl and Kay Curci Foundation, The Firmenich Foundation, and the Stanford MCHRI Transdisciplinary Initiatives Program (J.A.K.).

