## Supplementary Table 1 for "Spontaneous enteric nervous system activity precedes maturation of gastrointestinal motility"

**Fig.1H Clustering events/10 min**

| **ANOVA summary** | |
|---|---|
| F | 9.88 |
| P value | 0.0054 |
| P value summary | ** |
| Significant diff. among means | Yes |
| R squared | 0.6871 |

| **ANOVA table** | SS | DF | MS | F (DFn, DFc | P value |
|---|---|---|---|---|---|
| Treatment (between columns) | 3.163 | 2 | 1.582 | F (2, 9) = 9. | P=0.0054 |
| Residual (within columns) | 1.441 | 9 | 0.1601 | | |
| Total | 4.604 | 11 | | | |

| **Tukey's multiple comparisor** | Mean Diff. | 95.00% CI c | Summary | Adjusted P Value |
|---|---|---|---|---|
| E16.5 vs. E17.5 | -2.367 | -5.532 to 0.7 | ns | 0.1749 |
| E16.5 vs. E18.5 | -3.6 | -7.190 to -0. | * | 0.0492 |
| E17.5 vs. E18.5 | -1.233 | -4.399 to 1.9 | ns | 0.6107 |

**Fig.1I Minutes clustered/10 min**

| **ANOVA summary** | |
|---|---|
| F | 4.151 |
| P value | 0.0241 |
| P value summary | * |
| Significant diff. among means | Yes |
| R squared | 0.1917 |

| **ANOVA table** | SS | DF | MS | F (DFn, DFc | P value |
|---|---|---|---|---|---|
| Treatment (between columns) | 8.091 | 2 | 4.046 | F (2, 35) = 4 | P=0.0241 |
| Residual (within columns) | 34.11 | 35 | 0.9747 | | |
| Total | 42.21 | 37 | | | |

| **Tukey's multiple comparisor** | Mean Diff. | 95.00% CI c | Summary | Adjusted P Value |
|---|---|---|---|---|
| E16.5 vs. E17.5 | -0.7274 | -1.680 to 0.2 | ns | 0.1631 |
| E16.5 vs. E18.5 | -1.265 | -2.346 to -0. | * | 0.0187 |
| E17.5 vs. E18.5 | -0.538 | -1.491 to 0.4 | ns | 0.3612 |

**Fig.1H High energy events/10 min**

| **ANOVA summary** | |
|---|---|
| F | 1.409 |
| P value | 0.2821 |
| P value summary | ns |
| Significant diff. among means | No |
| R squared | 0.1901 |

| **ANOVA table** | SS | DF | MS | F (DFn, DFc | P value |
|---|---|---|---|---|---|
| Treatment (between columns) | 3.6 | 2 | 1.8 | F (2, 12) = 1 | P=0.2821 |
| Residual (within columns) | 15.33 | 12 | 1.278 | | |

| **Tukey's multiple comparisor** | Mean Diff. | 95.00% CI c | Summary | Adjusted P Value |
|---|---|---|---|---|
| E16.5 vs. E17.5 | -1 | -3.132 to 1.1 | ns | 0.4476 |
| E16.5 vs. E18.5 | -1 | -2.741 to 0.7 | ns | 0.3112 |
| E17.5 vs. E18.5 | 0 | -2.132 to 2.1 | ns | >0.9999 |

**Fig.2C Clustering events/10 min**

| **Multiple paired t tests** | Discovery? | P value | ean of Baseli | ean of 1uM T | Difference | E of differen | t ratio | df | q value |
|---|---|---|---|---|---|---|---|---|---|
| E16.5 | No | 0.078141 | 0.4286 | 0 | 0.4286 | 0.202 | 2.121 | 6 | 0.118383 |
| E17.5 | No | 0.177966 | 1.5 | 0 | 1.5 | 0.9574 | 1.567 | 5 | 0.179746 |
| E18.5 | No | 0.051946 | 2.333 | 0 | 2.333 | 0.9189 | 2.539 | 5 | 0.118383 |

**Fig.2D Minutes clustered/10 min**

| **Multiple paired t tests** | Discovery? | P value | ean of Baseli | Mean of TTX | Difference | E of differen | t ratio | df | q value |
|---|---|---|---|---|---|---|---|---|---|
| E16.5 | No | 0.090842 | 0.1847 | 0 | 0.1847 | 0.09178 | 2.012 | 6 | 0.13408 |
| E17.5 | No | 0.132753 | 0.3298 | 0 | 0.3298 | 0.1838 | 1.794 | 5 | 0.13408 |
| E18.5 | No | 0.040168 | 1.194 | 0 | 1.194 | 0.4336 | 2.753 | 5 | 0.121708 |

**Fig.2G High energy events/10 min**

| **Multiple paired t tests** | Discovery? | P value | ean of Baseli | ean of 1uM T | Difference | E of differen | t ratio | df | q value |
|---|---|---|---|---|---|---|---|---|---|
| E16.5 | No | 0.174688 | 0.3333 | 0 | 0.3333 | 0.2108 | 1.581 | 5 | 0.264652 |
| E17.5 | No | 0.269703 | 1.333 | 0 | 1.333 | 0.8819 | 1.512 | 2 | 0.2724 |
| E18.5 | No | 0.062352 | 1.333 | 0 | 1.333 | 0.5578 | 2.39 | 5 | 0.188928 |

**Fig.2J Clustering events/10 min**

| **Multiple unpaired t tests** | Discovery? | P value | lean of Contr | -Cre1;R26R | Difference | E of differen | t ratio | df | q value |
|---|---|---|---|---|---|---|---|---|---|
| E16.5 | Yes | 0.003605 | 3.143 | 0 | 3.143 | 0.8053 | 3.903 | 9 | 0.00182 |
| E17.5 | Yes | 0.002085 | 4.333 | 0 | 4.333 | 0.4303 | 10.07 | 3 | 0.00182 |
| E18.5 | No | 0.242871 | 1 | 0 | 1 | 0.8103 | 1.234 | 11 | 0.081767 |

**Fig.2K Minutes clustered/10 min**

| **Multiple unpaired t tests** | Discovery? | P value | lean of Contr | -Cre1;R26R | Difference | E of differen | t ratio | df | q value |
|---|---|---|---|---|---|---|---|---|---|
| E16.5 | Yes | 0.001596 | 1.376 | 0 | 1.376 | 0.309 | 4.452 | 9 | 0.003223 |
| E17.5 | No | 0.037039 | 2.715 | 0 | 2.715 | 0.7565 | 3.589 | 3 | 0.03741 |
| E18.5 | No | 0.225053 | 0.1862 | 0 | 0.1862 | 0.1449 | 1.285 | 11 | 0.151536 |
